# Hidden viral players: Diversity and ecological roles of viruses in groundwater microbiomes

**DOI:** 10.1101/2025.05.30.656956

**Authors:** Akbar Adjie Pratama, Olga Pérez-Carrascal, Matthew B. Sullivan, Kirsten Küsel

## Abstract

Groundwater ecosystems harbor diverse microbial communities adapted to energy-limited, light-deprived conditions, yet the role of viruses in these environments remains poorly understood. Here, we analyzed 1.26 terabases of metagenomic and metatranscriptomic data from seven wells in the Hainich Critical Zone Exploratory (CZE) to characterize groundwater viromes. We identified 257,252 viral operational taxonomic units (vOTUs) (≥5 kb), with 99% classified as novel, highlighting extensive uncharacterized viral diversity. Viruses exhibited a distinct host range, primarily targeting Proteobacteria, Candidate Phyla Radiation (CPR) bacteria, and DPANN archaea. Notably, CPR lineages displayed low virus-host ratios and viral CRISPR targeting multiple hosts, suggesting a “virus decoy” mechanism where they may absorb viral pressure, protecting bacteria hosts. Additionally, 3,378 vOTUs encoded auxiliary metabolic genes (AMGs) linked to carbon, nitrogen, and sulfur cycling, with viruses targeting 31.5% of host metabolic modules. These findings demonstrate viruses’ influence on microbial metabolic reprogramming and nutrient cycling in groundwater, shaping subsurface biogeochemistry.

## INTRODUCTION

Viruses that infect bacteria (bacteriophages/phages) are the most abundant biological entities on Earth, with an estimated population of 10^^31^, vastly outnumbering their microbial hosts^1,2^. Their ubiquity across diverse ecosystems highlights their adaptability and ecological significance.

In marine environments alone, the concentration of virus-like particles (VLPs) is estimated at 10^^30^, ranging from 10^^6^ to 10^^8^ VLPs per milliliter^3^. Research in marine systems has underscored their critical role in global biogeochemical cycles, particularly through mechanisms such as the ‘viral shunt’—which recycles organic matter within surface waters—and the ‘viral shuttle,’ which enhances microbial aggregation and contributes to carbon export via the ocean carbon biological pump^1–5^. Furthermore, ocean viruses drive microbial evolution and ecosystem functioning by facilitating the daily transfer of 10^^29^ genes^6^, including those encoding metabolic proteins such as photosynthetic reaction centres^7^. Given that viruses infect 20-40% of marine microbes at any given time^8^, they not only regulate host populations via lysis but also reprogram microbial metabolism and influence community dynamics via auxiliary metabolic genes (AMGs)^5,9,10^. These processes profoundly affect microbial evolution, nutrient cycling, and ecosystem stability.

Despite extensive research on viral ecology in marine and surface freshwater environments such as rivers^11^, and lakes^12^, the diversity, function, and ecological impact in subsurface aquatic ecosystems remain largely unknown. Groundwater, Earth’s largest reservoir of unfrozen freshwater, sustains ecosystems and plays a crucial role in global biogeochemical cycles, including carbon^13–15^, nitrogen^16,17^, sulfur^18,19^, phosphorus^20^, and various metals^21^. It is a microbially dominated habitat, characterized by nutrient scarcity, low organic carbon availability, and the absence of light, which collectively impose strong selective pressures on resident microbial communities^22^. Chemolithoautotrophy, a primary mode of carbon fixation in these environments, occurs at rates comparable to those in sunlit oligotrophic oceanic waters^13^, enabling microbial persistence in these extreme settings. Notably, groundwater harbors an abundance of ultra-small prokaryotes, including members of the Candidate Phyla Radiation (CPR) and DPANN superphylum archaea^23–25^, which possess highly streamlined genomes and are thought to rely on episymbiotic or syntrophic interactions for survival^22,23,26,27^. These microorganisms, which can constitute more than 50% of the microbial community in some groundwater systems^28^, likely play fundamental roles in subsurface biogeochemical cycling.

However, the role of viruses in shaping groundwater microbial communities remains largely unexplored. Their influence on key microbial groups, such as chemolithoautotrophs and ultra-small prokaryotes, as well as their influence on microbial adaptations to the nutrient-limited subsurface, remains unknown. This gap is particularly critical in the context of climate change, as declining groundwater levels due to rising temperatures and shifting precipitation patterns are expected to impact microbial dynamics and biogeochemical processes^29,30^. Understanding viral activity in these systems is essential for predicting how microbial interactions and nutrient cycles may respond to environmental change.

Here, we examined viral ecogenomic across seven groundwater wells within the Hainich Critical Zone Exploratory (CZE) in Germany. This well-characterized model system spans a range of physicochemical gradients, from oxic to anoxic conditions^31^, and hosts diverse microbial populations, including under-studied chemolithoautotrophs (e.g., *Nitrospiria* and *Sulfurifustaceae*) and ultra-small prokaryotes (e.g., CPR bacteria and DPANN archaea)^23–25^. We hypothesize that viruses modulate groundwater microbiomes by selectively infecting and shaping the abundance of these specialized microbial groups. Given the oligotrophic and hypoxic nature of these pristine ecosystems, we further propose that groundwater harbors novel, highly adapted viral lineages with distinct ecological strategies. By elucidating the diversity and functional roles of viruses in groundwater, this study provides a first step toward understanding viral contributions to subsurface microbial evolution and ecosystem functioning in one of Earth’s most expansive yet least explored habitats.

## RESULTS AND DISCUSSIONS

### Groundwater virus communities are diverse, novel, and unique

Our dataset is derived from a previous study of the Hainich CZE, which consisted of 0.515 terabases of short-read metagenomic data from 31 groundwater samples sampled from six wells spanning diverse hydrochemical and geochemical conditions (**Fig. 1a–c**). To these, we added 0.725 terabases of metagenomic data from 34 samples. These new samples were collected three years apart from the same six wells as the previous study, plus one additional well, resulting in a total of seven wells. This combined dataset totals 1.26 terabases across 65 samples. To improve the recovery of species-level viral operational taxonomic units^32^ (vOTUs), we also added six long-read metagenomes. Additionally, we leveraged ∼22 gigabases of metatranscriptomic data from 17 groundwater samples collected in 2015^19^.

**Figure 1.**
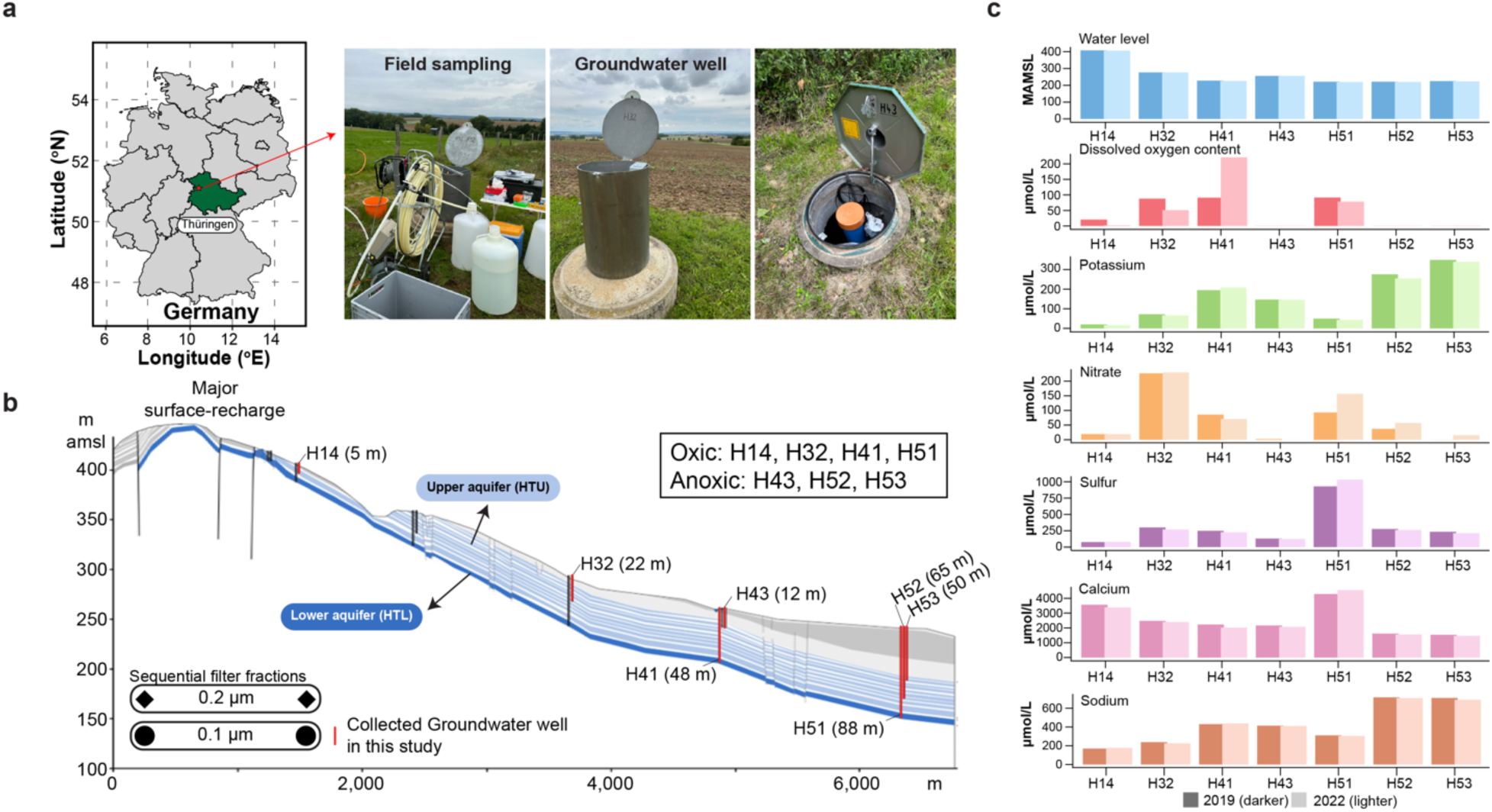
Groundwater sampling design. (**a**) Sampling was done in 2019 and 2022 in the Hainich Critical Zone Exploratory (CZE) located in Thuringia, Germany (red star in the map to the left). Photos show field sampling equipment and close-ups of one of the groundwater wells. (**b**) Schematic cross-section of the Hainich CZE sampling site. Sampled groundwater wells (vertical red lines) are located at different depths located in a hillslope area. The groundwater monitoring transects span ∼6 kilometres. The multistorey aquifer has a lower, karstified main aquifer (blue color, labelled HTL for Hainich transect lower aquifer) and an upper mudstone-dominated aquifer (teal color, labelled HTU for Hainich transect upper aquifer). Groundwater sampling was done via sequential filtration through 0.2-(diamond) and 0.1-µm (circle) pore size filters, with the numbers representing sampling replicates (the figure is modified from Kohlhepp *et al*., 2017; Hermann *et al.,* 2019). (**c**) Selected hydrochemical parameters of groundwater of the seven wells. Water level values for H43 were taken from the subsequent two measurement campaigns, as there were no measurements available at the time of the metagenome sampling. Other parameters can be seen in **Table S7**.

We first identified virus contigs and clustered these into viral operational taxonomic units (vOTUs), using a threshold of ≥ 95% average nucleotide identity over 80% of the smallest contig^33^. Because our datasets were cellular-fraction metagenomes, we applied additional filtering steps to remove potential microbial sequences in our final set of vOTUs. This included the removal of vOTUs with potential genes found in genomic islands, (**Figs. S1, S2**, see materials and methods). In total, 4,708,626 virus contigs from a total of 23,996,780 assembled contigs (≥ 1 kb) were identified (19.62%), generated from the 65 short-read and six long-read metagenomes (**Table S1**). These ∼4.7M virus contigs were further clustered into 257,252 (≥ 5 kb) vOTUs, 82,245 of which were ≥10 kb (**Table S2**). The evaluation of sampling saturation revealed accumulation curves that were nearing but not likely saturated (**Fig. S3a**), indicating high virus richness and under-sampling without virus-focused sequencing.

**Figure 2.**
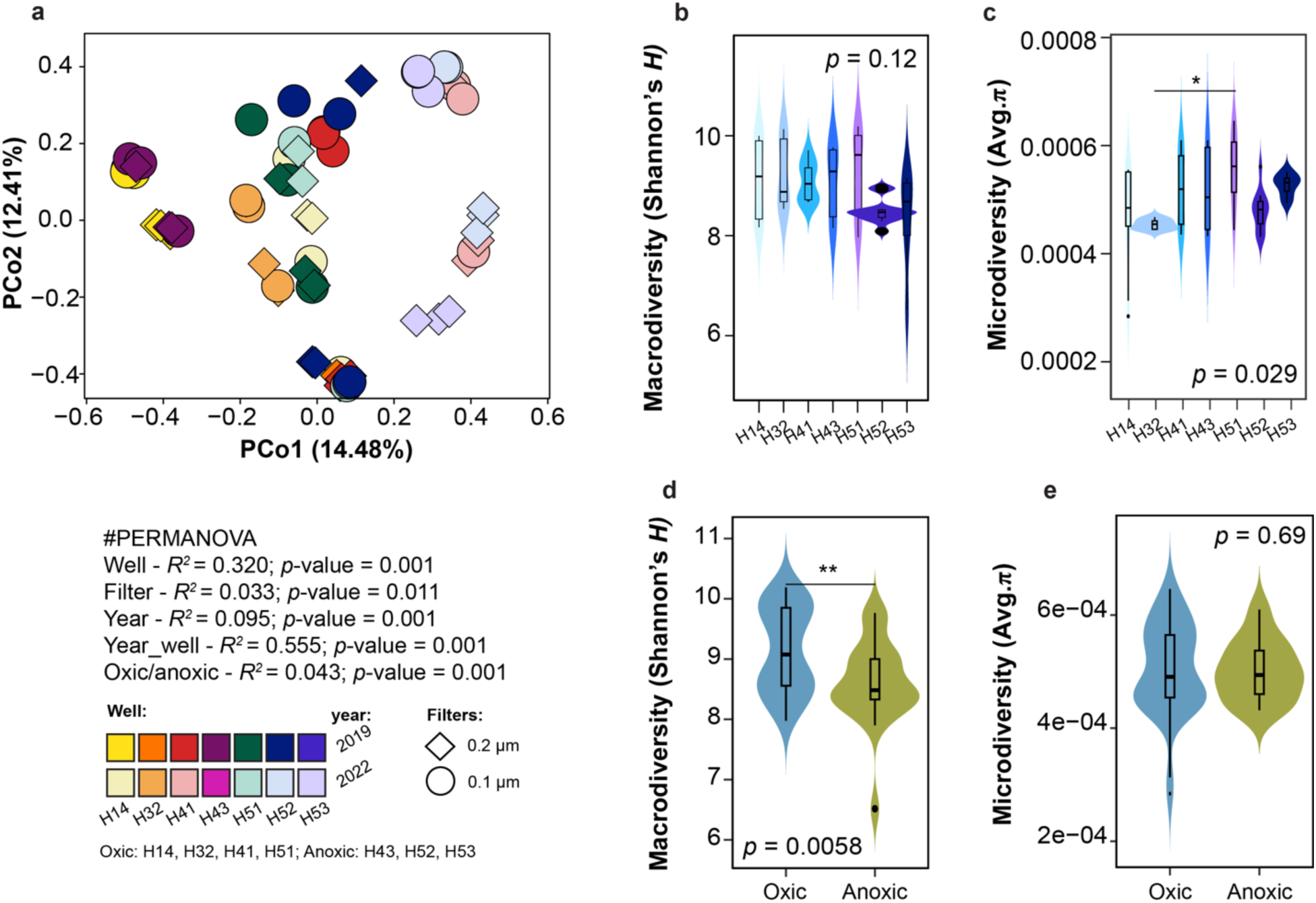
Virus community patterns along the groundwater monitoring transect. (**a**) Principal coordinate analysis (PCoA) of the groundwater viruses’ community-level Bray-Curtis dissimilarities was used to assess community structure. These data were derived from the relative abundance of 257,252 vOTUs (≥ 5 kb), where each point plotted represents this metric for one of the 65 sampled metagenomes color-coded by the well of origin. PERMANOVA was used to test for significant differences in viral community composition between wells, based on Bray-Curtis dissimilarity. (**b-c**) Macrodiversity (Shannon’s *H*) Microdiversity (Average π) diversity across the wells. All pairwise comparisons were shown, and statistically significant (\**p*<0.05) analyses were conducted using the Kruskal-Wallis test. (**d-e**) Macrodiversity (Shannon’s *H*) Microdiversity (Average π) across oxic and anoxic wells. The comparison between oxic/anoxic systems was shown, and statistically significant (\*\**p*<0.01) analyses were done using the Mann-Whitney U test.

To assess the novelty of the groundwater viruses, we used a hierarchical taxonomy approach (via vConTACT3 gene-sharing networks Bolduc *et al.,* under review), which revealed that 71.66% (*n* = 58,936) of vOTUs could be assigned to virus phylum. Of these, at the class-level, 99.42% (*n* = 58,593) were classified as Caudoviricetes, while 0.58% were classified as Laserviricetes, Tokiviricetes, Faserviricetes, and Vidaverviricetes. At the order-level, 13% belonged to a novel order of Caudoviricetes, Laserviricetes, and Tokiviricetes; 86.76% belonged to the order Methanobavirales and Kalamavirales; and 0.23% belonged to the unassigned order of Faserviricetes and Vidaverviricetes. At the family-level, 99.52% of the sequences belonged to the families Methanobavirales, Caudoviricetes, Kalamavirales, Laserviricetes, and Tokiviricetes; 0.24% belong to the family Zobellviridae, Mesyanzhinovviridae, Fredfastierviridae, Peduoviridae, Suoliviridae, Casjensviridae, Helgolandviridae, and Stanwilliamsviridae. The remaining 0.23% of the sequences could not be affiliated with any known family and are, therefore, unassigned to a particular taxonomic category. Finally, at the genus level, 99.75% of the sequences belong to novel genera, while 0.25% remain unassigned (see **Figures S4** and **Table S3**).

We next asked how many vOTUs were detectably active within the metatranscriptomes (*n* = 17), using previously established approaches and cut-offs^13,19^—where viruses are considered “active” if at least one expressed gene was detected per 10 kb of genome^34^ (see materials and methods). Application to our groundwater datasets revealed that 23.57% (*n* = 19,383) of vOTUs met this criterion (**Fig. S5a**), with significant variation in activity between wells (*p*-value < 0.0001). Well H41 had the highest relative activity, representing ∼25% of total transcript expression (see **Table S4**). To put this result into perspective, ∼58% of vOTUs were identified as active in permafrost soils, using a similar approach^34,35^, and up to 73% in Arctic soils using stable isotope probing^36^. In groundwater, active viruses were linked to diverse microbial hosts spanning 52 phyla (**Fig. S6a**), with Proteobacteria and Nitrospirota showing the highest associations across the wells (**Fig. S6b**).

The level of viral diversity observed in our study (∼82K vOTUs ≥10 kb) substantially expands virus diversity estimates in groundwater ecosystems. Prior to this analysis, the available number of groundwater vOTUs in IMG/VR^37^ (an aggregative database) was 3,580 - a 23-fold increase. Specifically, groundwater virus-focused studies in New Zealand^38^ and the southeastern coast of Sweden^39^ had found 1,571 (52.7-fold over) and 1,407 vOTUs (59-fold), respectively. Thus, by the number of vOTUs alone, our study stands out among the scarce data available for groundwater viruses. Relative to other aquatic biomes, this represents almost half (∼0.4-fold) of the total vOTUs found in global ocean studies (Global Ocean Virus Database, version 2, *n* = 195,728)^40^ and far exceeds the number reported for the deep oceanic trench (*n* = 1,628)^41^, and rivers (*n* = 1,230)^11^. Notably, our vOTU count is 2.8-fold lower (*n* = 230,944) than the current estimate of the GOV dataset^42^.

To further assess novelty, we compared these groundwater vOTUs (only those of ≥ 10 kb, *n* = 82,756; see materials and methods) and their protein clusters (PCs)—groups of proteins clustered based on sequence similarity that represent shared functional or evolutionary traits—to those available in public databases, including IMG/VR v4.1^37^, Gut Virome Database (GVD)^43^, Global Soil Virome database (GSV)^44^, Global Oceans Viromes 2.0 database (GOV2)^40^, Viral RefSeq v.222^45^, Fennoscandian Groundwater Virome (FGV)^39^, and Rumen Virome Database (RVD)^46^ (**Fig. S7**). At the vOTU level, not a single sequence from any of these datasets clustered with our new groundwater viruses. This suggests strong groundwater endemism, given the lack of shared taxa at the species level. However, PC-level similarities revealed some overlaps (average of 3.25%), consistently at only a few percent. The highest similarity was observed with GSV^44^ at 7.32% (*n* = 62,898 PCs), followed by IMG/VR^37^ groundwater at 4.43% (*n* = 14,152 PCs) and the global ocean virome (GOV2)^40^ at 4.38% (*n* = 30,895 PCs). The percentage of PCs shared with other databases was even lower at <2% (**Fig. S7**). Broadening our search, we queried against freshwater vOTUs from IMG/VR and found only one VOTU in common (from rivers), and no VOTUs were shared with viruses from lentic, reservoir, lotic, pond, stream ecosystems, or other sampled groundwater systems (**Fig. S8a-b**).

These results highlight the strong endemism of groundwater viruses, with minimal overlap in vOTUs or encoded PCs or genus-level virus clusters that were similar to sequences in various publicly available groundwater-related databases. This result is consistent with findings from comprehensive metagenomic studies showing that only 7% of novel metagenomic protein families are shared across ecosystems, highlighting the uniqueness and limited cross-biome distribution^47^. Consistent with this, analysis of the groundwater microbial communities reveals a notable proportion of unique taxa specific to individual wells, suggesting localized novelty and potential endemism within the groundwater system^28,48,26^.

### Ecological patterns of the virus community are well-specific

Previous studies on microbial communities in these wells, based on 16S rRNA sequencing and metagenome-assembled genomes (MAGs), revealed substantial variation between wells^28,48^. We hypothesized that virus communities would exhibit similar well-specific patterns. To test this, we assessed vOTU abundances (via metagenomic read mapping) across all 65 samples from the 7 wells (**Fig. S9a**), and organized vOTUs in two categories (*i*) unique (one well only) or shared (multiple wells) vOTUs, and (*ii*) core (vOTUs present in more than 50% of the wells), common (present in 21–50% of the wells), and rare (present in 1–20% of the wells) vOTUs^49,50^. This revealed that (*i*) 58.23% of vOTUs were unique, 39.73% were shared, and 2.04% vOTUs were not categorized. While in (*ii*) 6.10% were identified as a core, 33.63% were common, and 58.23% were rare vOTUs across all wells, and 2.04% vOTUs were not categorized (**Fig. S9b**). We also found the highest percentage (19.15%) of unique vOTUs in the uphill shallow well (H14), while shared vOTUs were more common in wells at the bottom of the hillslope (H52 and H53). These wells also had a higher percentage of core and common vOTUs (**Fig. S9b-c**). To align our findings with previous work, we compared vOTU distribution and host predictions to bacterial patterns from a decade-long 16S rRNA study^51^. That study showed distinct community structures, with wells H52 and H53 dominated by core bacteria (55–65%) largely classified as Parcubacteria (CPR lineage), while other wells had lower core fractions (9–29%) but higher diversity (Shannon’s *H* 5.2–5.9). Consistently, our host predictions identified vOTUs likely infecting Paceibacteria (a class within Patescibacteria, the MAG-based equivalent of Parcubacteria), comprising 1.98% and 1.28% of vOTUs in H52 and H53, respectively, supporting a link between viral and microbiome community structure. These differences likely result from the contrasting hydrological connectivity of the upper and lower aquifers in the Hainich CZE (see **Fig. 1b**). Wells H52 and H53 access the hydrologically isolated upper aquifer, where alternating carbonate (limestone) and siliciclastic (marlstone) layers restrict water exchange. In contrast, the lower aquifer, accessed by more hydrologically connected wells (H14, H32, H41, H51), facilitates greater water movement^31^. This hydrological contrast is mirrored in the geochemical conditions: connected wells exhibit oxic conditions, while isolated wells range from suboxic to anoxic^51^. Notably, oxic wells harbored ∼3.3-fold more unique vOTUs than anoxic wells (**Fig. S9a**). This pattern is not exclusive to groundwater systems; a similar trend has been observed in marine environments, where viral diversity is ∼2.5-fold higher in well-oxygenated waters compared to oxygen minimum zones (OMZs)^52^.

Next, we assessed virus community composition patterns through spatial and temporal lenses and hypothesized that abiotic factors likely play an indirect role through host dynamics. To assess these patterns, we examined whether viral communities correlated with microbial shift gradients across wells, and whether temporal variation (sampling year) also played a role. Using non-metric multidimensional scaling (NMDS) and principal coordinates analysis (PCoA) revealed a clear virus community structure (**Fig. 2a** and **Table S5**). Specifically, ordination revealed that the combined effect of well and year (year_well) was the strongest driver of viral community structure (PERMANOVA, R² = 0.555, *p* < 0.001). In contrast, individual factors such as well location, sampling year, filter size, and oxic/anoxic conditions had weaker influences on viral composition (**Fig. S10a**). These findings suggest that spatial and temporal variation within specific wells plays a key role in shaping viral community dynamics.

We next examined how macrodiversity (inter-population diversity) and microdiversity (intra-population genetic variation) changed along the groundwater monitoring transect (**Figs. S10-S13** and **Table S6**). Macrodiversity (Shannon’s *H*) did not vary significantly between the wells (*p* = 0.12, Fig. 2b), nor across time (Wilcox test, *p* <0.01; 2 years of sampling) despite the variations in Shannon’s *H* diversity. These results further supported the consistency of the virus diversity over time (**Fig. S11**). However, a significant difference (*p* <0.01) in macrodiversity was observed between oxic and anoxic wells (**Fig. 2d-e**). This likely reflects the strong influence of oxygen availability on potential host cells, as oxic conditions support higher microbial diversity (**Fig. S14-S15**; see in particular **Fig. S15i** with Wilcox test, *p* <0.01) and activity (2.25-fold higher; **Fig. S16b**). Increased microbial activity and diversity may lead to greater virus diversity through an increase in host availability. Microdiversity, as assessed through average nucleotide diversity (π), varied significantly between wells (**Fig. 2c**) highlighting the fine-scale genetic differentiation within viral populations. One possible explanation for these patterns lies in the contrasting roles of macro- and microdiversity in shaping viral community dynamics. While macrodiversity, reflecting the overall taxonomic composition of the viral community, may remain relatively stable across wells due to consistent nutrient inputs or hydrological patterns, microdiversity captures genetic variation within individual viral populations and is more sensitive to localized ecological processes. Specifically, microdiversity is shaped by mechanisms such as isolation-by-environment, where hydrological barriers or geochemical gradients (see **Fig. 1C**) restrict viral dispersal and promote localized genetic differentiation^40,53^. Additionally, host-virus interactions play a critical role, as viral replication, mutation rates, and horizontal gene transfer (HGT) vary depending on host availability and selective pressures within each well (see virus-host ratio later in this section). These localized processes can increase genetic variability within viral populations— manifesting as high microdiversity—even when the broader community composition (macrodiversity) remains unchanged^54^. This distinction underscores how viral microdiversity serves as a sensitive indicator of fine-scale ecological dynamics, revealing how environmental heterogeneity and host-virus co-evolution shape genetic variation in groundwater viromes.

### Ecological drivers of the groundwater viruses

To further test whether environmental factors drive virus richness, macrodiversity, microdiversity, as well as host richness and diversity, we analyzed 18 different environmental variables (**Table S7;** see materials and methods). Mantel analysis revealed that temperature significantly correlated with virus richness (Mantel’s *p* = 0.01–0.05), while ammonium and potassium concentrations (Mantel’s *p* <0.01) were significantly correlated with virus macrodiversity (**Fig. S17**). Host richness and Shannon’s H were significantly correlated with water level, pH, redox potential, base neutralizing capacity (acidity), and magnesium concentrations (**Fig. S17**).

While the local geochemical factors were strongly associated with shaping the overall virus community composition (macrodiversity), none were found to significantly impact microdiversity. Moreover, these factors may drive virus macrodiversity by influencing viral replication rates, host availability, and selective pressures. For example, potassium and ammonium have previously been linked to virus community structure and activity, including transcriptional activity, dissolved organic carbon (DOC) concentrations, and iron cycling in groundwater^38^. These factors may indirectly shape viral macrodiversity by altering host physiology, metabolic potential, and susceptibility to infection. Furthermore, studies of soil virome using a similar Mantel analysis have consistently shown that temperature, nutrients (e.g., ammonium, potassium), and pH influence microbial and viral communities’ abundance^53^. While specific drivers may vary across ecosystems, geochemical factors have been shown to influence virus and microbial communities, including within groundwater systems.

### Groundwater viruses are predicted to infect diverse prokaryotic phyla

Understanding viral host range is essential for elucidating their role in shaping microbial community dynamics and ecosystem functions. To identify potential hosts for groundwater viruses, we employed an *in silico* approach that integrates multiple sequence-based features from both viral and cellular genomes and 1,275 metagenome-assembled genomes (MAGs) to predict host-virus associations (see materials and methods). We found (at high or ≥ 90% host confidence score) that 11.62% (*n* = 9,615) of vOTUs were associated with hosts MAGs spanning 60 phyla, 143 classes, 339 orders, and 650 families (**Fig. S18** and **Table S8**). An unexpected result in this analysis was that vOTUs were predominantly associated with the least abundant microbial taxa, i.e., Proteobacteria as the most common host (22.9% of vOTU links) (**Fig. 3a-b**, **Fig. S18b**). This pattern aligns with other groundwater virome studies, which have also reported a strong association with Proteobacteria^38^. However, it contrasts with marine ecosystems, where viruses typically infect the most abundant taxa, such as SAR11^55^. This divergence suggests that viral-host interactions in groundwater may be shaped by additional factors beyond host abundance, such as metabolic activity, niche specialization, and hydrological constraints. To further explore this relationship, we used the metatranscriptomic data from our MAGs, which revealed that Proteobacteria were one of the most active taxa in the groundwater (**Fig. S16**). These results suggest that viral targeting is shaped by host activity and metabolic capacity rather than abundance alone. Although Patescibacteria were also highly active, their reduced genomes may limit virus-host interactions, while metabolically versatile Proteobacteria were more frequently linked to vOTUs.

**Figure 3.**
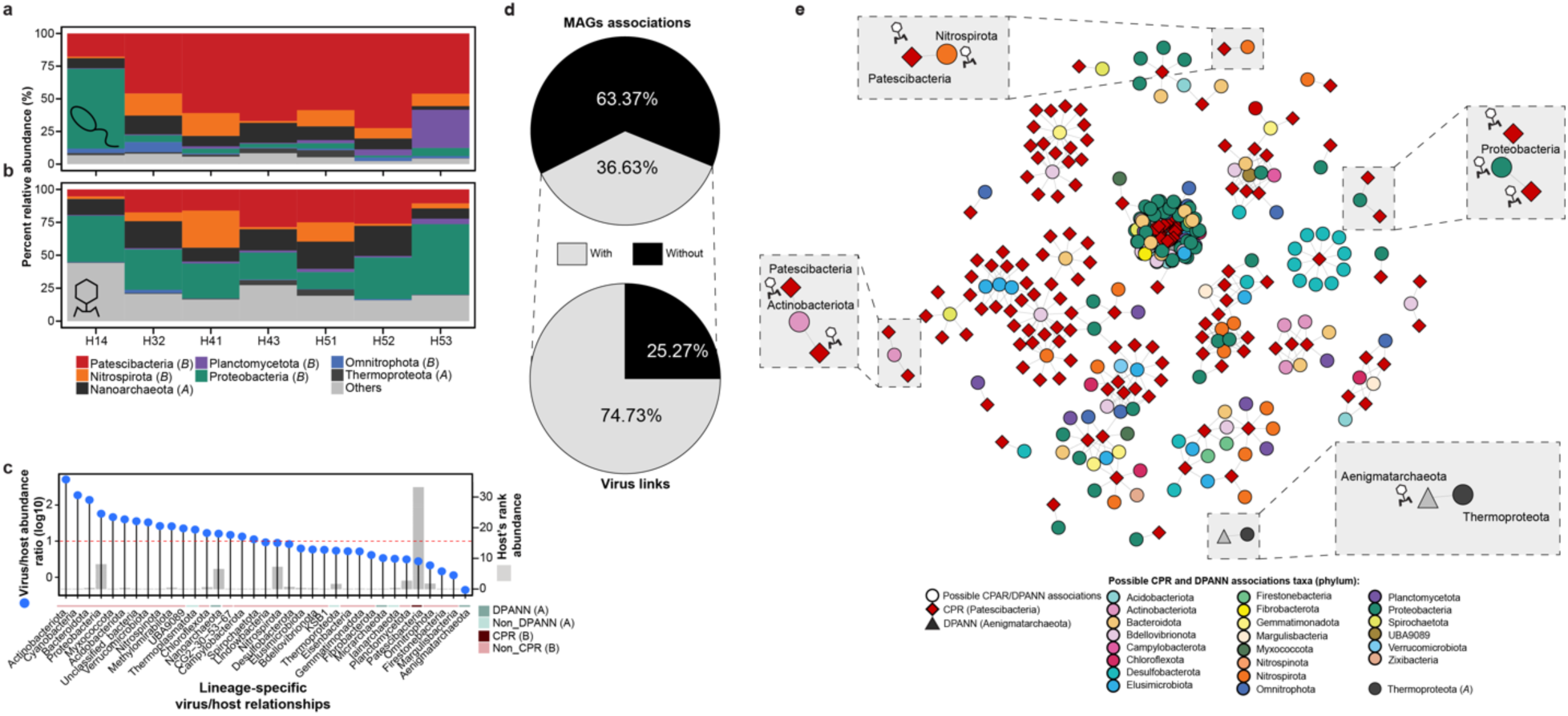
Virus operational taxonomic unit-level analysis of *in silico* host prediction and tripartite CPR/DPANN-host-virus relationships. (**a**) Host relative abundance (%) for each sample (*n* = 65) for the predicted host. The colors indicate the phylum, and others represent all additional phyla observed that were < 1 % of the total relative abundance. *(A)* for archaea and (*B*) for bacteria. (**b**) Virus relative abundance (%), colored by the predicted hosts’ phylum. (**c**) Lollipop plots depict the virus/host abundance ratios by host lineage, calculated as the ratio of per base-pair average coverage depth from read mapping to viral contigs and host population genomes, respectively, normalized by the number of sequencings reads in each sample. The dots indicate the mean VHR across the 65 metagenomes, and the horizontal red dashed line indicates the 1:1 virus: host abundance ratio. The bar plots depict the host’s rank abundance (the number of MAG per taxa can be seen in Table SX). The line colours are the categories for DPANN, non-DPANN, CPR, and non-CPR. (**d**) Pie charts illustrate the overall number of MAGs associated with other bacterial and archaeal MAGs (top) and, among these, the proportion linked to viruses (bottom). (**e**) Co-occurrence network to assess the potential association of CPR and DPANN with other bacterial and archaeal MAGs for 2019 sampling year. The co-occurrence analysis for 2022 sampling year can be seen in **Fig. S18**. The circles depict the potential bacteria hosts of CPR and DPANN. The red diamonds represent CPR, and the dark-grey triangle is for DPANN. The zoomed-in clusters highlight some associations of CPR and DPANN MAGs with potential host MAGs, e.g. Chloroflexota, and Proteobacteria. Virus links in the inserted figure were identified through *in silico* host prediction analysis. Pie charts illustrate the percentage of MAG (*n* = 1,275) with CPR/DPANN associations based on the co-occurrence network analysis.

To quantify viral pressure across microbial lineages, we calculated lineage-specific virus/host abundance ratios (VHRs), a proxy for viral pressure on microbial populations^34,35^. VHRs were calculated as the ratio of per-base-pair average coverage depth from read mapping to viral contigs and host population genomes, respectively, adjusted for the total number of sequencing reads in each sample^34^. Among the 35 lineages analyzed, 48.57% (17 out of 35) had VHRs greater than 1 (log10 scale), indicating that viral populations outnumbered their respective hosts. The highest VHRs were observed in Actinobacteria, Cyanobacteria, Bacteroidota, and Proteobacteria (**Fig. 3c**; **Table S9**) suggesting strong viral control over these microbial groups. These VHRs varied across wells (**Table S9**), reflecting environmental heterogeneity (**Fig. S19-S20**). In oxic wells (H14, H32, H41, H51), Proteobacteria showed elevated VHRs, consistent with higher viral diversity in oxygen-rich environments that support active microbial growth. Conversely, Patescibacteria showed higher VHRs in anoxic wells, likely driven by distinct viral-host interactions adapted to low-oxygen conditions. These lineage-specific patterns are consistent with host metabolic strategies: Proteobacteria thrive in nutrient-rich, oxic conditions, whereas Patescibacteria specialize in anoxic environments. Similar observations in Omnitrophota from anoxic permafrost soils highlight how high VHRs can enhance viral lysis, thereby influencing carbon turnover and biogeochemical cycles^34,35^.

Despite being one of the most abundant microbial groups (59.39% relative abundance), Patescibacteria exhibited a VHR below one, indicating that viral populations do not outnumber their hosts as they do for other microbial taxa. This is particularly intriguing given that *in silico* host analysis linked 18.42% of vOTUs (*n* = 1,771) to Patescibacteria, suggesting extensive viral interaction with this group. In addition, our metatranscriptomic data revealed a high number of viruses linked to Patescibacteria (23.56%, see **Fig. S6a**), suggesting that CPR frequently encounters viral interactions.

One possible explanation for these patterns is that Patescibacteria and other CPR bacteria act as ‘virus decoys,’^56^ attracting viral interactions away from their hosts, and without supporting productive infections. Alternatively, the low VHRs could reflect frequent lysogenic infections (see next section) or physical protection through symbiotic associations^23^ that shield them from active viral predation. Additionally, Patescibacteria may possess uncharacterized antiviral defense mechanisms, contributing to the low ratio of viral to host abundance despite frequent encounters suggested by host prediction and transcriptional activity. If CPR bacteria limit viral replication, they may divert viral pressure away from other prokaryotes, effectively influencing host-virus dynamics at a community level. To test this hypothesis, we investigated CRISPR-Cas spacer (i.e., proto-spacer) matches between viruses and both CPR/DPANN and non-CPR/non-DPANN bacterial and archaeal MAGs. We searched for and identified CRISPR-Cas spacers, then aligned them using BLAST (*e*-value = 1e-05; ≥95% identity, ≥90% coverage, allowing one-zero mismatches) to assess potential matches against 82,245 vOTUs (≥10 kb). This analysis revealed two proto-spacers (2 vOTUs) matching Desulfobacteriota and Patescibacteria (Paceibacteria), and seven proto-spacers (7 vOTUs) matching Nitrospirota and Nanoarchaeota (Nanoarchaeia) (**Table S10**).

Interestingly, Patescibacteria (Paceibacteria) are known to retain the standard bacterial genetic code (code 11), but some subgroups may have alternative genetic codes that have not been fully characterized. A similar result has been reported CRISPR-Cas spacer matches identified in 23 viruses targeted by both CPR (Gracilibacteria) and non-CPR bacteria, with some lacking in-frame TGA codons, allowing replication in both host types^57^. A separate groundwater study predicted seven CPR viruses infecting non-CPR phyla, including a Gracilibacteria virus capable of replicating in both Gracilibacteria (code 25) and Bacteriota (code 11) using compatible genetic code mechanisms^58^. These results suggest that CPR/DPANN may act as viral bait, attracting viral interactions that could otherwise impact their associated hosts. This potential buffering role highlights how CPR/DPANN taxa may influence viral-host dynamics in groundwater, shaping ecosystem stability by modulating viral pressures across different microbial lineages.

### Tripartite relationship among episymbiotic prokaryotes, their host and viruses

Groundwater ecosystems are known to harbor small cell-sized bacteria and archaea (e.g. CPR and DPANN^23–25,28,48^), and often form close physical associations with other microbes. In addition to the viruses targeting CPR and non-CPR bacteria^57,58^, interactions between Patescibacteria, their methylotrophic proteobacterial hosts, and jumbo phages have also been observed in freshwater samples, forming a tripartite interaction^59^. Building on this concept, we aimed to assess potential tripartite interactions by using (*i*) *in silico* host prediction analysis combined with co-occurrence analysis to assess CPR/DPANN-host associations and (*ii*) prophage identification in CPR/DPANN MAGs. First, to explore virus-host linkages for CPR and DPANN at both ecosystem-wide and class-level scales, we conducted *in silico* virus-host prediction analysis^60^ (**Fig. S21**). Across all sampled groundwater wells, 2.14% of vOTUs (*n* = 1,771) were linked to CPR, and 2.17% (*n* = 1,796) to DPANN (**Fig. S21a-c**). Within Patescibacteria (CPR), Paceibacteria had the highest number of associations (*n* = 1,001), followed by Microgenomatia (*n* = 343), ABY1 (*n* = 209), and Gracilibacteria (*n* = 126). For DPANN, Nanoarchaea showed the most connections (*n* = 1,759) (**Fig. S21d-e**). Previous groundwater virome studies^38^ have reported 18 vOTUs links to CPR and one vOTU link to Nanoarchaeia. In a separate study^61^, 97 non-redundant DPANN viruses were linked to 8 DPANN phyla. Collectively, these findings represent a significant increase in virus-host linkages compared to previous studies of groundwater viruses.

To further investigate potential host associations for CPR and DPANN, we performed a co-occurrence network analysis using a normalized relative abundance matrix. To mitigate the limitations of correlation-based inference, we performed the co-occurrence analysis separately for each year to account for potential temporal variability and to provide a clearer picture of meaningful associations specific to each period, rather than averaging across multiple years^62^ (**Fig 3d-e**, and **Fig. S22**). Overall, the analysis revealed that 36.62% (467/1,275) of MAGs were associated with other MAGs (**Fig. 3d**), of which 212 CPR MAGs were associated with non-CPR bacteria and 14 DPANN MAGs were associated with other non-DPANN archaea. Within CPR, Patescibacteria exhibited associations with microbial taxa such as Chloroflexota, Desulfobacterota, Nitrospirota, Proteobacteria, and Omnitrophota, while Nanoarchaeota showed associations with Iainarchaeota (**Fig. 3d-e**; **Table S11**). These associations have been previously reported^25,48,63^, although only a few have been cultured and experimentally verified, such as CPR– Pseudomonadota (Halochromatium)^64^ and DPANN–Thermoproteota (Metallosphaera)^65^. Of the MAGs with detected associations, 74.73% (349/467) were also linked to viruses. Of these, 159 were CPR (FCPU426 and Patescibacteria) MAGs, and 13 were DPANN (Aenigmatarchaeota, Iainarchaeota, and Nanoarchaeota) MAGs. These results highlight the dynamic interactions between potential episymbiotic microbes, their hosts, and viruses, reinforcing the hypothesis that CPR/DPANN taxa experience frequent viral encounters that may influence their ecological roles in groundwater.

Lastly, we assessed the presence of prophages in CPR/DPANN MAGs to further investigate viral interactions within these microbial communities. The detection of prophages in CPR/DPANN provides evidence of viral integration, revealing possible past or ongoing interactions between CPR/DPANN, their non-CPR/non-DPANN hosts, and associated viruses. Among the 599 prophage vOTUs (≥5 kb) identified from 23.53% (300/1,275) of MAGs, we found prophages in 20.42% (106/519) of Patescibacteria MAGs and 29% (38/131) of Nanoarchaeota MAGs (**Fig. S23).** While prophages are common among bacteria, their presence in host-dependent CPR/DPANN suggests viral integration may play a functional role in their interactions with host microbes. Overall, these findings expand our understanding of the role of viruses in shaping microbial consortia and highlight the complex interplay between episymbiotic prokaryotes, their hosts, and viruses in groundwater ecosystems.

### Viruses’ influence on biogeochemical processes in groundwater

To further evaluate the ecological role of viruses in groundwater ecosystems, we examined vOTUs for the presence of auxiliary metabolic genes (AMGs). These genes are randomly obtained from host genomes, but if they offer a selective advantage to the virus (presumably by enhancing host processes during virus infection), they can become fixed in the virus genomes^34,66,67^. To find AMGs, we adapted a recently proposed approach^68^ that leverages automated functional annotation using DRAM-v processing^69^ and applied it to 82,245 (≥10 kb) vOTUs (see materials and methods). This revealed that 4.11% (*n* = 3,378) of vOTUs encoded 4,093 putative AMGs (**Fig. 4a-b**), which were grouped into 377 protein clusters (**Table S12**). Functionally, these AMGs were involved in transport and central carbon, amino acid, nitrogen, and sulfur metabolism (**Fig. S24**). Interestingly, when we compared the AMGs found in this study (*n* = 4,093) with those found in the global ocean^42^, we found 75.6% overlap in AMGs, while 58.83% of the groundwater AMG were also found with those found in the 20-years of freshwater lake virome study^70^ (**Fig. S24b**). Furthermore, AMG-carrying viruses were found to be associated with a wide taxonomic range of hosts, which spanned 24 phyla, 44 classes, 75 orders, and 122 families (**Table S12**), hinting that AMG acquisition as a common virus strategy to enhance their reproduction.

**Figure 4.**
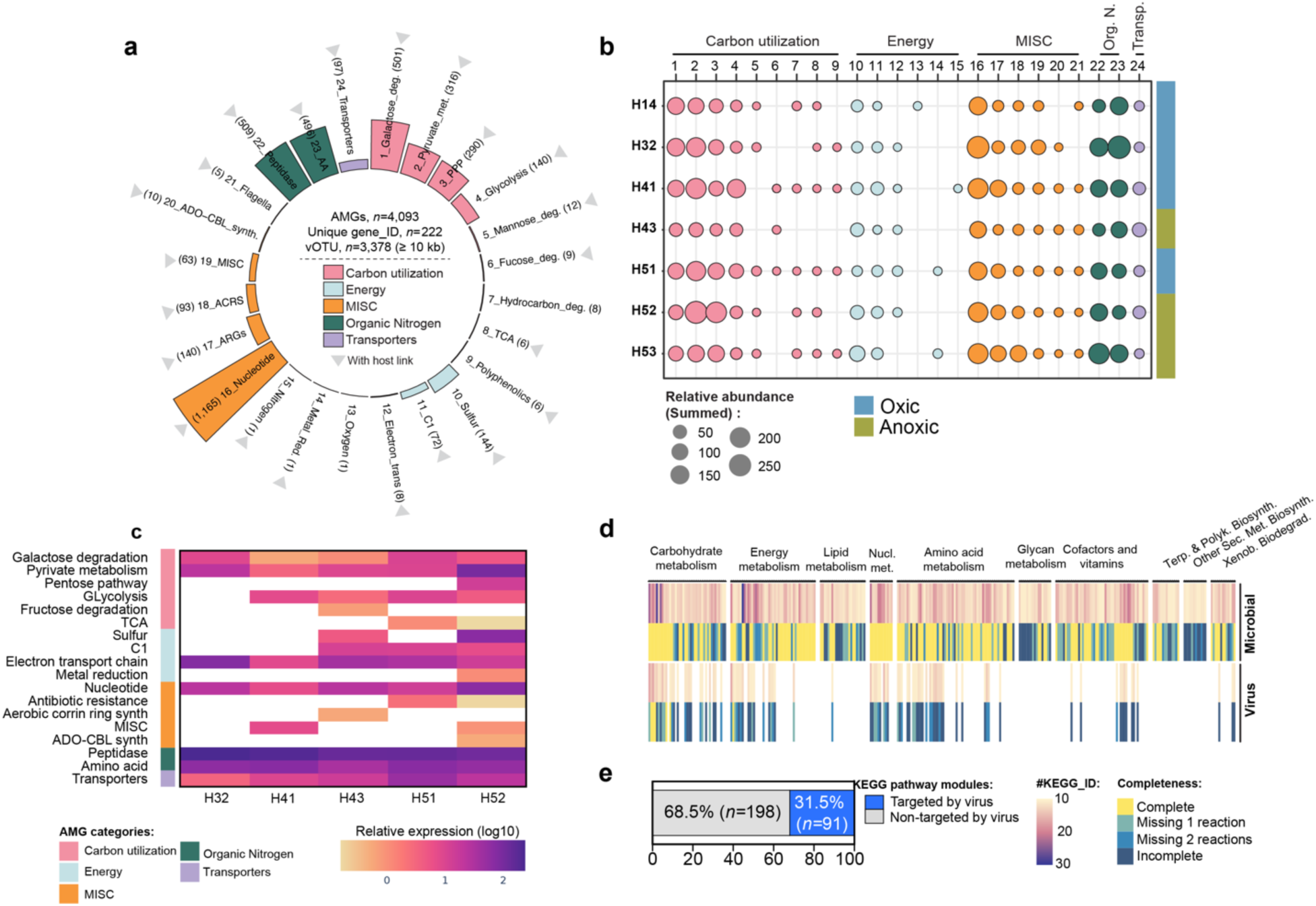
The distribution and metatranscriptomics analysis of the putative auxiliary metabolic genes (AMGs) **(a)** circular bar plot depicts the identified AMGs. The colors represent the category of the AMGs. The grey triangles indicate the host links were found. **(b)** The bubble plot depicts the distribution of viruses-carrying AMGs per category across the well. The relative abundance is represented in the size of the bubble, and the oxic-anoxic well is indicated by blue-sky and olive-green, respectively. **(c)** The expressions of the AMG (relative expression in log10). The colour intensity on the heatmap reflects the relative expression level of each virus within a specific metatranscriptome sample. Higher TPM corresponds to more intense colours. **(d)** Heatmap represents the reconstruction of microbial metabolic pathways for microbial (top) and putative AMGs (bottom). The heatmap colors represent the number of KEGG IDs, and the completeness of the metabolic pathways is indicated. Terp. & polyk. Biosynth. stands for terpenoid and polyketide biosynthesis, Sec.Met. Biosynth. stands for secondary metabolite biosynthesis, and Xenob.biodegrad. stands for Xenobiotic Biodegradation. The detailed metabolic reconstruction can be seen in **Table S13**. **(e)** The stacked bar plot depicts the percentage of KEGG pathway modules targeted (in blue), and non-targeted by viruses (in grey).

Given the oligotrophic nature of groundwater, where resource recycling and utilization efficiency are paramount, we hypothesized that auxiliary metabolic gene (AMG) occurrence and activity would be particularly high. Indeed, we identified 8.04% (*n* = 686) of expressed genes as AMGs (**Fig. 4c**), associated with key metabolic pathways including the TCA cycle, glycolysis, and nitrogen and sulfur metabolism, highlighting the potential role of viruses in shaping microbial energy flow in these ecosystems. Notably, transcript levels of putative AMGs increased along the well gradient, with downstream wells exhibiting higher expression of AMGs related to peptidase activity (∼45.6% relative abundance), amino acid metabolism (∼14.37%), pyruvate metabolism (∼9.29%), and the electron transport chain (∼9%) (**Fig. 4c**). This pattern suggests that as microbial and viral communities stabilized across wells, with a higher fraction of core taxa, viruses played an increasing role in host metabolic processes. These findings align with previous studies showing that AMGs involved in central carbon metabolism are particularly important in nutrient-limited environments, such as the deep ocean^71^, where they redirect host energy production to enhance viral replication. This underscores the crucial role of viral AMGs in optimizing energy flow and resource utilization in oligotrophic environments.

To assess this in more detail, we focused on the AMGs relevant to key biogeochemical cycles in the groundwater ecosystem (**Fig. S24-S25**). First, we evaluated sulfur cycling AMGs, including *cysA, cysH*, and *cysE*, which host prediction analysis suggested were linked to viruses infecting Proteobacteria, Nitrospirota, and Desulfobacterota – all key taxa known for sulfate reduction and assimilation previously reported in Hainich groundwater^16,19^. Viruses carrying sulfur-related AMGs, which are critical for sulfur fixation and the transformation of organosulfur compounds, have been found in various domains and microbiomes across the globe and appear to be more abundant than previously thought^72^. Second, nitrogen-related AMGs, such as *nrfA*, which facilitates nitrite reduction to ammonia, were associated with Bdellovibrionota and only found in well H41 (**Fig. 4b**), a site with the highest nitrification rates^17^. Third, AMGs targeting central carbon and methane-related metabolism were also observed. Genes such as *cofE*, involved in coenzyme F420 biosynthesis, highlight the potential for viruses to regulate methane flux, while AMGs related to the TCA cycle and reductive Wood-Ljungdahl pathway were associated with Bacteroidota, potentially enhancing microbial carbon fixation^13^. Although groundwater contributes a small fraction (0.2%) of global methane emissions^73^, the presence of methane-related AMGs suggests that viruses may play an important role in regulating methane cycling in these ecosystems.

Taken together, these results provide compelling evidence that groundwater viruses actively shape microbial metabolism, influence nutrient cycling, and drive ecosystem adaptation in resource-limited subsurface environments.

### AMGs in CPR bacteria and DPANN archaea

To investigate the potential influence of AMGs on prevalent microbial groups within this ecosystem, we focused on AMGs found in vOTUs predicted to infect CPR bacteria and DPANN archaea. Our results showed that 7.1% (*n* = 125) of CPR vOTUs carried AMGs related to carbon metabolisms (e.g., TCA cycle, pentose phosphate pathway, glycolysis), amino acid metabolism, and nutrient transporters (**Fig. S26**; **Table S12)**. Similarly, 5.8% (*n* = 105) of DPANN viruses carried AMGs associated with carbon utilization (e.g., glycolysis, pyruvate metabolism), energy production (e.g., sulfur metabolism), and organic nitrogen metabolism (**Table S12**). Notable AMGs, such as pyruvate dehydrogenase and threonine/serine dehydratases, may enhance host metabolic activity alongside chaperone and cell surface-related genes^59^. While these AMGs could indirectly influence episymbiotic interactions by affecting host energy production and nutrient recycling, it is crucial to emphasize that the primary function of viral AMGs is to optimize host metabolism for viral replication^74^. The observed AMGs likely serve to redirect host resources to favor viral propagation, and any effects on episymbiotic relationships would be secondary and potentially incidental

The prevalence of AMGs related to carbon utilization and energy production suggests that viral infections may reshape host metabolism to prioritize resource acquisition and viral replication in resource-limited groundwater environments. In such oligotrophic settings, hosts may possess limited internal building blocks, necessitating viral acquisition of additional resources to facilitate viral assembly. The increased presence of AMGs within the viral population could reflect this heightened demand for scavenging available nutrients. This pattern mirrors findings from aphotic marine ecosystems, where viromes from deep-sea environments have shown an enrichment of AMGs involved in central carbon metabolism^71^. In marine systems, these genes have been proposed to redirect host metabolism via the glyoxylate shunt, increasing energy production while potentially suppressing amino acid synthesis. Such a metabolic shift maximizes efficiency under extreme nutrient limitations, allowing for viral replication and in some cases, potentially enabling host survival through modulated metabolic activity. However, it’s important to note that viral replication typically leads to host cell lysis. Instances of persistent infections, where viruses ‘bud out’ of the host without killing it, have been observed in some archaeal viruses^75^, providing a potential exception. The presence of viral AMGs in both groundwater and marine systems suggests a convergent ecological strategy, where viruses modulate host metabolism to optimize resource acquisition and replication in extreme environments. These findings highlight the broader role of viruses in shaping microbial ecosystem function beyond marine environments, particularly in deep, oligotrophic ecosystems where microbial communities persist under energy-limited conditions.

To provide an overview of the detectable microbial metabolisms that are present in these groundwater wells and to assess the extent to which viruses reprogram these host metabolisms, we mapped community-level MAG-derived metabolic enzymes against KEGG ortholog (KOs) and then overlaid this with identified AMGs (see materials and methods). This revealed that viruses target 31.5% (*n* = 91) of host metabolic modules, for example, Citrate cycle (M00009), and Cobalamin biosynthesis (M00122) (**Fig. 4d-e**, **Table S13**), highlighting their role in shaping microbial metabolisms. While a global study of marine viruses reported a higher prevalence of AMGs (19% of virus populations, with ∼9% observed)^42^, viruses in groundwater and marine ecosystems also target comparable host metabolic pathways—31.5% in groundwater and 37.6% in oceans^42^—suggesting the ecological importance of AMGs are likely important in microbial reprogramming across diverse environments.

Taken together, these findings highlight the central role of AMGs in driving nutrient cycling in subsurface ecosystems and may enhance microbial metabolic adaptation. By enabling microbial hosts to optimize metabolic pathways under nutrient-limited and oxygen-variable conditions, viruses play a critical role in maintaining groundwater ecosystem functioning. Integrating these viral mechanisms into models of groundwater biogeochemistry could improve our understanding of ecosystem resilience and nutrient flux dynamics in response to environmental change.

### Conclusion

This study reveals the critical yet understudied role of viruses in shaping the groundwater microbiome and driving biogeochemical processes in subsurface ecosystems. We uncovered a vast and largely uncharacterized viral diversity, with ∼99% representing novel genera, including a higher-than-expected prevalence of viruses associated with small-genome bacteria and archaea, particularly CPR and DPANN episymbionts. These findings suggest that viral interactions within the groundwater microbiome are more abundant and extensive than previously recognized, with potential consequences for microbial adaptation and ecosystem function. These viruses carry auxiliary metabolic genes (AMGs) linked to carbon, nitrogen, and sulfur cycles, highlighting their role in modulating host metabolism and influencing nutrient cycling in energy-limited environments. Our findings provide a foundational framework for understanding virus-driven microbial interactions in the groundwater microbiome, emphasizing their influence on host metabolic reprogramming and ecosystem stability.

While *in silico* analyses have identified potential hosts^19,28,48^more high-resolution approaches, such as Hi-C^76^, epic-PCR^77^, and recent integrated single-molecule DNA fluorescence in situ hybridization (FISH)-highly multiplexed ribosomal (r)RNA-FISH^78^ approaches are needed to validate virus-host interactions. Additionally, the lack of cultured representatives for CPR/DPANN and their hosts presents a major challenge in characterizing infection dynamics and evolutionary relationships. Cultivating these organisms and their associated viruses will be essential for resolving key mechanistic questions, including how viruses infect ultra-small prokaryotes, evade host defenses, and exploit host machinery. Future research should develop experimental model systems to investigate groundwater-specific virus-host interactions, ultimately refining our understanding of viral contributions to microbial evolution, adaptation, and biogeochemical cycling in the groundwater microbiome.

## MATERIALS AND METHODS

### Sample collection, DNA extraction, and metagenomic sequencing

Pristine groundwater metagenomes from samples collected in the Hainich Critical Zone Exploratory (CZE) in Thüringen, Germany, during January 2019 (*n* = 31) as previously described, and new metagenomes collected in November 2022 (*n* = 34) (**Table S1**). A total of 50 and 100 L of groundwater were filtered from each well through 0.2 μm filters and 0.1 μm filters per triplicate, and DNA extractions were performed to the captured cells using a phenol-chloroform method. A total of 65 metagenomes (total read = 8,737,844,712; range = 134,996,654-100,861,616; Q1 = 112,553,890.5; and Q3 = 153,430,689.5) were sequenced using the Illumina NextSeq 500 system and paired-end library (2 × 150 bp), while for 2022 metagenomes the NovaSeq 6000 system was used. Characteristics and sampling details are described elsewhere^28,31,48^. Briefly, in the characteristics and sampling details, there are two distinct well groupings based on location and oxygen availability. Wells H41 and H51 (oxic) are connected to the main aquifer (Limestone-rich Upper Muschelkalk) and situated in flatter areas. In contrast, wells H14 and H32 (oxic) are found in uphill zones, while wells H43, H52, and H53 (suboxic or anoxic) are located downhill (**Fig.1**).

A previous study demonstrated that long-read sequencing substantially enhances the recovery of vOTUs^32^. To build on the short-read sequencing data, we incorporated subsamples from long-read sequencing (*n* = 6), selecting those with the highest DNA concentrations for long-read metagenomic analysis. Nanopore libraries were prepared using approximately 500 ng of metagenomic DNA and the ligation kit (SQK-LSK110, ONT). The detailed numbers of reads and bases can be seen in **Table S1**.

### Metagenomics processing, identification, and abundance quantification of MAGs

The metagenomics-assembled genomes (MAGs) were generated as previously mentioned^48^. Briefly, metagenomic sequencing was quality-filtered via the bbduk script (BBMap version 39.01). Next, SPAdes (version 3.15.2) (--meta mode and other parameters are default)^79^ was used to assemble the contigs, and contigs longer than 1,000 bp were binned using five binning tools, using default settings including (*i*) binsanity (version 0.2.7)^80^ with 5 kbp as a minimum scaffold size, (*ii*) abawaca (version 1.0.0) using 5 kbp, and minimum scaffold size, (*iii*) maxbin2 (version 2.2.6)^81^ using 40 and 107 gene markers, (*iv*) CONCOCT (version 1.0.0) and (*v*) metabat2 (version 2.12.1)^82^. Metawrap (version 1.3.2) refinement (filters -c 50 -x 10)^83^ was then carried out to refine bins obtained from the five binning algorithms and obtain the best representative MAGs using dRep (version 3.4.0)^84^. We also used *anvi-script-gen-CPR-classifier* script from Anvi’o v6.1^85^, which uses a supervised machine learning model (random forest classifier) for predicting the probability of the MAGs to confirm the CPR genomes (https://merenlab.org/2016/04/17/predicting-CPR-Genomes/). From the initial set of 1,778 dereplicated metagenome-assembled genomes (MAGs) of high and medium quality, further refinement was performed using established thresholds, including: (*i*) a quality score (QS = completeness – 5 × contamination), with bins scoring QS < 50 being removed^86^; (*ii*) retention of high-quality bins with completeness ≥ 90% and contamination ≤ 5%; and (*iii*) retention of medium-quality bins with completeness > 50% and contamination ≤ 10%^87^. A genome-quality assessment was carried out using the CheckM workflow (version 1.1.3)^88^. Lastly, the taxonomic annotations of the MAGs selected for analysis were carried out with GTDB-Tk (version 1.5.1)^89^ using GTDB (release 202) as a reference database. Following this refinement process, a total of 1,275 MAGs were retained for subsequent *in silico* virus-host analysis.

For long-read processing, the samples were then individually sequenced on MinION flow cells (FLO-MIN106 with an R9.4.1 pore) for 72 hours. The basecalling was performed using the Guppy software (Version 6.0.1) with the super accurate model (dna_r9.4.1_450bps_sup.cfg). Next, the fastq reads were used to generate hybrid assemblies with Spades Hybrid. A total of 17 hybrid assemblies were generated for the samples belonging to the filter fraction 0.2 um) (see **Fig. S1**).

To create a MAG abundance table, we calculated trimmed mean abundances (-m trimmed_mean) using CoverM --min-read-percent-identity 0.95 --min-read-aligned-percent 0.75 --min-covered-fraction 0.25) (https://github.com/wwood/CoverM). Next, we normalized the trimmed mean abundance table by dividing the abundances by the number of quality-controlled reads per sample per 1 Tb^90^. Approximately 25.9% of metagenomic reads mapped to the MAGs, indicating a substantial proportion of the microbial community is represented by the reconstructed genomes. The accumulation curves for these MAGs demonstrated overall saturation across the groundwater metagenomes, suggesting that most of the microbial diversity captured by MAGs has been adequately sampled (**Fig. S12c**).

### Virus identification, filtering, virus clustering, and quality assessment

Viromics pipeline can be seen in **Fig S1**. For virus identification, in addition to assembled contigs generated from metaSPAdes as described above, we also used assembled contigs generated from MEGAHIT (default)^91^. Further, we utilized VirSorter2 (version 2.2.3)^92^, DeepVirFinder (version 1.0)^93^, VIBRANT (version 1.2.1)^94^, and geNomad (version 1.5.1)^95^ to detect viruses within assembled contigs from each sample. To ensure the confidence of the identified viruses, we applied the following filters, the first one follows the recommended thresholds for the individual tool, including, (*i*) for VirSorter2 we applied standard operational procedure (protocols.io/view/viral-sequence-identification-sop-with-virsorter2-5qpvoyqebg4o/v3). (*ii*) for DeepVirFinder, the score was ≥ 0.9, and the *e*-value was ≤ 0.05. (*iii*) default settings for VIBRANT, and (*iv*) geNomad. Given the conservative nature of CheckV in identifying virus and host genes can serve as a valuable tool for identifying false positives^96^. This aligns with prior research that has explored the necessity of additional filtration steps aimed at removing non-viral sequences^68^. Hence, we applied a second filter, utilizing CheckV, recommended from previous studies^92,97^, and manual spot check for DeepVirFinder, VIBRANT, and geNomad. For this, we retain virus contigs with (*i*) virus genes > 0, (*ii*) virus gene = 0 and host gene = 0, (*iii*) virus contigs length ≥ 1 kb, and iv) those with unknown genes ≥ 75%^97^. Lastly, (*iv*) we also trimmed the host regions using the CheckV “end-to-end” function and default settings (see detailed workflow and documented numbers of virus contigs **Fig. 1**). To ensure high confidence, we also employed manual spot checks (**Fig. S2**). The resulting 4,708,626 virus contigs were clustered into a viral operational taxonomic unit (vOTU) at 95% ANI across 80% of alignment fraction (relative to the shorter sequence)^33^ using MMSeq2^98^ resulting in 2,412,499 (≥ 1 kb), of which 257,814 (≥ 5 kb), and 82,807 (≥ 10 kb) vOTUs.

To further remove contigs that are more likely degraded signals in cellular hypervariable genomic islands or more likely to be giant viruses or mobile elements, we adopted a curation rule filter^42^ to vOTUs ≥ 100 kb in length size. In this filter, we exclude any contigs that contained genes annotated as transposonses, lipopolysaccharide genes (glycosyltransferase, nucleotidyl transferase, carbohydrate kinases, nucleotide sugar epimerase), endonuclease, integrase, or plasmid stability since such genes are enriched in the genomic island (reviews see Dobrindt *et al*.^99^ and Bertelli *et al*.^100^). This removed 562 vOTUs > 100 kb such that our final dataset of conservatively identified vOTUs was 257,252 (≥ 5 kb), of which 82,245 (≥ 10 kb) vOTUs. We further used CheckV^96^ to assess the quality and completeness of vOTUs, and VIBRANT^94^ to predict virus lifestyle (virulence vs temperate, both using default settings) (**Table S2**).

### Calculating virus abundance and ecological statistics

To create a virus abundance table, we calculated trimmed mean abundances (-m trimmed_mean) using CoverM (--min-read-percent-identity 0.95 --min-read-aligned-percent 0.75 --min-covered-fraction 0.7) (https://github.com/wwood/CoverM). Next, we normalized the trimmed mean abundance table by dividing the abundances by the number of quality-controlled reads per sample per 1 Tb (**Table S4**). We further assess species accumulation curves using the vegan specaccum() function in R, providing insights into the accumulation of distinct viral species across samples (**Fig. 1g**).

Further, we examined the distribution of each vOTU across the groundwater. We considered viruses to be present if the normalized vOTUs relative abundance exceeded 0 in at least one sample. Utilizing a similar approach, we further categorized vOTUs across the sample into “unique” viruses as vOTU found solely in one sample, and “shared” viruses were identified in multiple samples^40^. Moreover, we also categorized the vOTUs into, “core” (vOTUs present in more than 50% of the wells), “common” (present in 21–50% of the wells), and “rare” (present in 1–20% of the wells)^49,50^. To visualize the relationships and similarities of the vOTUs across the well, we utilized an UpSet plot generated using the UpSetR (**Fig. S3, S5**), and stacked bar plots using ggplot2 packages in R (**Fig. 1h, 1i,** and **Fig. S6**). Moreover, to investigate the similarity distribution of viruses at a finer level, focusing on genotypes, and to account for the potential of convergent evolution, we implemented a re-clustering approach. Specifically, we re-clustered the virus contigs at thresholds of ≥ 99.5%, ≥ 99.8%, and ≥ 100% of average nucleotide identity. Subsequently, we conducted similar analyses and visualized the results using UpSetR package in R (**Fig. S6**).

The ecological analysis was performed using the relative abundance table using R with the ‘vegan’ package. First, we log-transformed the normalized relative abundance table from the previous section and generated Bray-Curtis dissimilarity (β-diversity) matrices as input for Principal Coordinate Analysis (PCoA, cmdscale() function) (**Fig. S7**) and Non-metric MultiDimensional Scaling analysis (NMDS, metaMDS() function) (**Fig. 2a**, and **S7**). For PCoA, the emerging groups were statistically verified using PERMANOVA tests with adonis() function. For NMDS, statistical significance was determined using the anosim() function and multiple response permutation procedure (mrpp() function, distance = euclidean, permutations = 9999). We further calculated α-diversity metrics (Shannon’s *H*, Richness, Pielou’s J evenness, Simpson, Inverse-Simpson, and evenness) for each sample using the R package ‘vegan’. In the main text, we referred to Shannon’s *H* as a macrodiversity (inter-population or species diversity) (**Table S13**).

Pearson’s correlation and mantel test were performed to assess the pairwise relationships between the environmental factor data (*n* = 18), viruses (Shannon’s *H*, macro-, and microdiversity), and their respective hosts (Shannon’s *H*, and richness) (**Fig. 2e**). The physicochemical data that were used (**Table S6**), including, water level (MAMSL), water temperature (°C), specific electrical conductivity (µS/cm), pH, redox potential (mV), dissolved oxygen content (µmol/L), acid neutralizing capacity (Alkalinity) (meq/L), base neutralizing capacity (acidity) (meq/L), total inorganic carbon (µmol/L), chloride (µmol/L), calcium (µmol/L), potassium (µmol/L), magnesium (µmol/L), sodium (µmol/L), sulfur (µmol/L), nitrate (µmol/L), ammonium (µmol/L), sulfate (µmol/L)^51^. All analyses and data visualization were conducted using the linkET package (https://github.com/Hy4m/linkET) in R.

Virus microdiversity was calculated as previously described^40^, using MetaPop^101^ (--min_cov 70). Briefly, Viral populations meeting the criteria of an average read depth of ≥ 10x across 70% of their contig in at least one sample were analyzed for microdiversity. Single nucleotide variants (SNVs) were called from BAM files with reads mapping at ≥ 95% nucleotide identity, using a quality threshold of > 30. Nucleotide diversity (π) per genome was calculated following downsampling to 10x coverage per locus. The final microdiversity value (average π) for each sample was determined by averaging π values from 100 randomly selected viral populations across 1,000 subsamplings (**Fig. S15 –** Avg. π).

### Virus taxonomy classification

There is a wide range of tools available for virus taxonomy. We utilize the recent virus taxonomy tool vConTACT3 (https://bitbucket.org/MAVERICLab/vcontact3/src/master/) for all the vOTUs ≥ 10kb. The vConTACT3 classifies viruses into taxa based on the gene-network approach and is aligned with the International Committee on Taxonomy of Viruses (ICTV) taxonomy. The vConTACT3 uses NCBI Virus RefSeq release 220 and incorporates information from the Virus-Host DB (using GenBank release 257). We used the vConTACT3 default setting to assign the taxonomy of the vOTUs. The taxonomy assignment can be seen in **Fig. S4,** and the complete results can be found in **Table S3**.

### Viral clustering and database comparison

To compare the similarity of Hainich groundwater viruses with other publicly available databases, we downloaded IMG/VR v4.1^37^ (*n* = > 5M vOTUs), Gut Virome Database (GVD, *n* = 33,242)^43^, Global Soil Virome database (GSV, *n* = 80,750)^44^, Global Oceans Viromes 2.0 database (GOV2, *n* = 195,728)^40^, Viral RefSeq v.222^45^ (*n* = 18,719), Fennoscandian Groundwater Virome (FGV, *n* = 4,051)^39^, and Rumen Virome Database (RVD, *n* = 397,180)^46^. To ensure an equal comparison, we further filtered vOTUs ≥ 10 kb in length size. This process resulted in a refined dataset GVD (*n* = 15,330), GSV (*n* = 80,750), GOV2 (*n* = 195,728), Viral RefSeq v.222^45^ (*n* = 6,574), FGV (*n* = 1,407), and RVD (*n* = 193,350). For IMG/VR, we applied these categories, (*i*) vOTUs ≥ 10 kb in length size, (*ii*) only freshwater origin (lake, river, lentic, groundwater, creek, pond, lotic, and reservoir), (*iii*) with high-quality genome, (*iv*) exclude those assigned to RNA viruses and no taxonomy assignment. Totaling 558,408 IMG/VR freshwater databases.

In the first comparison analysis, we compared our datasets at the vOTU- and protein cluster-level against GOV2, GSV, GVD, virus RefSeq, RVD, and IMG/VR (groundwater only). For vOTU-level, we utilized three different approaches, first, we used blastn, as described previously^40^. The vOTUs with a nucleotide alignment at ≥ 95% nucleotide identity and an alignment length ≥ 50% the length was considered present in other databases (**Fig. 1d**). Additionally, we also compared vOTUs with other databases using MMSeq2 (--min-seq-id 0.95 -c 0.5 and --min-seq-id 0.95 -c 0.8). The results showed no overlapping. At the protein cluster level, amino acids clustered with other entries in the database were counted. The amino acid sequences were identified using Prodigal (version 2.6.1)^102^. Resulting in this study (*n* = 2,592,284), GOV2 (*n* = 5189693), GSV (*n* = 2,970,347), GVD (*n* = 636,701), IMG/VR ‘groundwater’ (*n* = 131,765), virus RefSeq (*n* = 654,189), FGV (*n* = 40,432), and RVD (*n* = 7,455,032). Moreover, we filtered and used ≥ 100 amino acids for further analysis, resulting in this study (*n* = 1,703,721), GOV2 (*n* = 3,618,709), GSV (*n* = 2,089,631), GVD (*n* = 446,574), IMG/VR groundwater (*n* = 95,888), virus RefSeq (*n* = 450,852), FGV (*n* = 25,811), and RVD (*n* = 5,023,585). To this end, we then clustered our dataset with other databases using MMSeq2 (parameters: --min-seq-id 0.3 -c 0.6 -s 7.5)^42^. Lastly, we compared our vOTUs with the IMG/VR freshwater database and GOV2 using MMSeq2 (--min-seq-id 0.95 -c 0.5 and --min-seq-id 0.95 - c 0.8) (**Fig. S3)**.

### Host predictions and lineage-specific virus/host abundance ratios (VHRs)

We utilized a recent host of virus identification tools, iPHop (version 1.3.2)^60^. iPHop integrated blast and CRISPR-spacer similarity with, oligonucleotide frequency (ONF) based distance/dissimilarity measures between a pair of DNA sequences, *k-*mer frequencies, the Gaussian Model (GM), and a machine-learning approach of protein cluster to assign hosts. For our analysis, briefly, we used two approaches, including default databases (iPHoP_db_Sept21_rw, Host genomes extracted from GTDB r202, IMG published genomes as of Sept. 2021, and GEMv1--Earth’s Microbiomes catalog) and 1,275 metagenomics assembled genomes (MAGs) (**Figs. S14** and **Table S7**) obtained from the same samples at our site. For the default, we used the ‘predict’ and ‘Sept_2021_pub_rw’ database. In the second approach, before the analysis, we followed the recommended workflow (https://bitbucket.org/srouxjgi/iphop/src/main/), we symbolically linked our 1,275 MAGs to the original database using ‘add_to_db’. To this end, we integrate the results from both approaches and only consider the top hits of virus-host links for further analysis. The VHRs were calculated as previously described^34^. In brief, by dividing the sum of coverage depth per base pair obtained from a read mapping of vOTUs and host population (MAGs), respectively. The lineage-specific virus/host abundance values can be seen in **Table S12**.

On the microbial side, of the 1,275 available Hainich MAGs, Patescibacteria dominates (59.39% relative abundance), followed by Nanoarchaeota (8.17%), Proteobacteria (6.98%), Thermoproteota (1.40%), and other taxa (<1%) (**Fig. S14**). Given that longer virome fragments (≥ 10 kb) improve host prediction accuracy by increasing recall rates and reducing false discovery rates (FDRs)^60^, we applied this threshold to our dataset. We combined cellular genomic data and predicted hosts for our 82,756 vOTUs (≥ 10 kb).

### CRISPR-Cas spacer identification and screening against viruses

To identify the CRISPR-Cas and spacers (≥3 direct repeats) from these MAGs, we used MinCED v0.4.2 (https://github.com/ctSkennerton/minced). Then matched against vOTUs (82,245; ≥10 kb) with ≤1 mismatch over ≥95% of the spacer length using BLASTn (-word_size 7 -task ‘blastn-short’ -evalue 1e-5). A two-mismatch of CRISPR-Cas spacers has been reported to represent an optimal tradeoff between capturing real phage-host interactions and minimizing false predictions^103^. I also matched the spacers against IMG/VR spacer databases (Virus DNA DB ver. 4 2022-09-15, Viral Spacers Public 2025-01-22, Metagenome Spacers Public 2025-01-22).

### Identification of virus-encoded auxiliary metabolic genes (AMGs)

To identify virus-encoded putative auxiliary metabolic genes (AMGs), we used DRAM-v^69^ and as recommended, we used ≥ 10 kb for this analysis. vOTUs DRAM-v utilized a variety of databases, such as Pfram, KEGG, UniProt, CAZyme, MEROPS (the peptide database), VOGDB (Virus Orthologous Groups Database), and NCBI viral RefSeq to annotate virus genomes. Prior, we performed VirSorter2 (--prep-for-drama) to generate the input files required for DRAM-v. We then applied several recommended filters to distill putative AMG, including (*i*) retained those with AMG category 1-3, and (*ii*) those with AMG flag M. Subsequently, those with AMG category 4 and with AMG flags V, A, P, and B were considered as not putative AMG. To ensure the confidence of putative AMGs (post-DRAMv curation), I refined the analysis by excluding specific viral genes commonly misclassified as metabolic genes by DRAM-v. For example, included genes for enzymes like dUTP pyrophosphatase^104^, DNA (cytosine-5-)-methyltransferase^105^, glycosyl transferase^106^, ribonucleoside-triphosphate reductase^107^, and ribosomal subunit genes. Additionally, we also annotate the microbial MAGs (*n* = 1,275) using DRAM^69^. We further clustered the putative AMGs using MMSeq2 (--cluster-mode 2 --cov-mode 1 -c 0.6 -s 7.5 --kmer-per-seq 20) as previously described^67^.

To delve deeper into the influence of putative AMGs in the metabolic pathways of groundwater microbial communities using the KEGG mapper construct^108^. Initially, microbial genes associated with KEGG Orthology (KOs) were mapped onto microbial metabolic pathways using KEGG mapper reconstruction. This step enabled the recording of two key parameters: the number of unique KEGG identities (no_kegg_ID) involved in module pathways, and an assessment of module completeness (module_comp). Concurrently, a similar analysis was conducted for putative AMGs-identified KEGGs, allowing for a comprehensive understanding of their role within these pathways. Subsequently, the obtained results were visually represented using ggplot2’s geom_tile() functions. Furthermore, we considered viruses targeted specific metabolic modules, by the presence of a putative AMG within those pathways. This integrated approach provided valuable insights into the intricate interplay between microbial and viral components within groundwater microbial communities and their metabolic activities.

### Metatranscriptome analysis

The metatranscriptomics data (initial total *n* = 18) and analysis were described elsewhere^13^. We excluded the metatranscriptomic data from well H42 because it did not correspond to the metagenomic data. Therefore, we used a total of 17 metatranscriptomic datasets. Briefly, the subsamples of groundwater metatranscriptomes were collected in August and November 2015, for wells H32, H41, H42, H43, H51, and H52. Since H42 was not used in this project, we then removed them from the analysis. The QAQC filtered reads were mapped to MAGs using Bowtie2 v.2.3.5 in sensitive mode^109^, and the total number of *rpoB* transcripts from each metatranscriptomic library was determined, as described in the preceding for metagenomes. Moreover, to quantify the abundances of transcripts from transcript data, the Kallisto package (version 0.48.0) was used^110^ and was normalized via scaling-factor calculations based on the total number of *rpoB* reads from the QAQC filtered metatranscriptomic reads. Additionally, for a viral population to be considered detected in a metatranscriptome, there must be an average coverage value for at least one gene for every ten kilobases (kb) of viral genomic sequence, as recommended previously^34^. These methods estimate ‘active’ viral community composition, but their accuracy remains uncertain due to the lack of standardized approaches or biological benchmarks for assessing viral ‘activity’ in bulk metatranscriptome data. We further visualized these active viruses using a heatmap from Flaski (https://github.com/mpg-age-bioinformatics/flaski)^111^ where log10 scaling was applied to the summed of transcripts relative expression (for viruses in **Table S16**). The scaling factor for *rpoB* can be seen in **Table S18.**

We also analyzed metatranscriptomic data for MAGs as described above. Genes were included if they were expressed in at least 10% of the samples, as previously described^112^. We also assess the relative expression without applying that cutoff as previously described^13^. In addition to scaling factor normalization, we further normalized relative gene expression using the average genome size of MAGs within each phylum. The average genome size was determined by estimating genome lengths from our MAGs (observed genome length/completeness) and integrating reference genomes from IMG/G (last accessed: 24 February 2025). This dataset includes genomes from prokaryotic isolates (115,873 bacterial and 1,379 archaeal genomes) and MAGs (14,481 bacterial and 1,002 archaeal genomes).

### Co-occurrence network analysis for MAGs

To identify the possible host for Candidate phyla radiation (CPR) we used a similar approach described previously elsewhere^48^. Initially, we utilized the normalized average genome coverages table encompassing 1,275 MAGs across all metagenomes. We then used this abundance matrix to calculate the proportionality of the coverage profiles in R package propR v5.0.2^113^. Further, we used the following thresholds: (*i*) *⍴* a threshold of 0.95 was used for network creation to highlight only the most relevant co-occurrences, (*ii*) only those that correlate with the CPR and DPANN, and (*iii*) those with positive correlations. The network was generated using Cytoscape v3.10.1^114^ for visualization.

## ACKNOWLEDGMENTS

This project is funded by the Deutsche Forschungsgemeinschaft (DFG, German Research Foundation) under Germanýs Excellence Strategy – EXC 2051 – Project-ID 390713860 and via the Collaborative Research Centre AquaDiva (CRC 1076 AquaDiva Project-ID 218627073) of the Friedrich-Schiller-University Jena.

Part of this work was enabled by funding provided to M.B.S by the U.S. Department of Energy, award #DE-SC0023307.

The authors would like to thank Heiko Minkmar, Falko Gutmann, René Maskos, and Stefan Riedel for groundwater sampling and on-site measurements/sample preparation. The authors would also like to thank Olivier Zablocki for his input on the manuscript.

## AUTHOR CONTRIBUTIONS

A.A.P., K.K. and M.B.S. created the study design. O.P.C., and A.A.P., collected all datasets. O.P.C., and A.A.P., performed the data analysis and visualization. A.A.P., O.P.C., M.B.S., and K.K. contributed to the scientific discussion and wrote the manuscript. All authors read and approved the final manuscript.

## SOM FIGURE AND TABLE LEGENDS

**Fig. S1. Bioinformatics workflow.**

Flow diagrams showing the bioinformatic workflow for metagenomic data preprocessing and virus metagenomics analysis, including virus identification and virus operational taxonomic units (vOTUs) analysis (light blue background), the metagenomic assembled genomes (MAGs) generation (lavender), the long-read NanoPore sequencing processing (lavender), viruses’ comparison against public databases at the vOTU- and Protein-cluster-level (light peach), an ecological analysis, including read-mapping, relative abundance table generation, α-, β-diversity, vOTU-distribution, and microdiversity analysis (light green), taxonomy analysis (light orange), potential host prediction analysis (light pinkish-orange), and depicts functional analysis for auxiliary metabolic genes (AMGs) analysis (light-yellow).

**Fig.S2. Example of contigs that were retained and removed after filtering steps.** The genomes are colored to illustrate microbial (green), viral (blue), and non-annotated genes (grey).

**Fig.S3. Overview of the groundwater vOTUs.**

**(a)** An accumulation curve of virus populations in the groundwater metagenomics (*n* = 65). The insert plot depicts the individual wells. The means are represented by coloured lines, and from randomizations of samples. **(b)** The groundwater 257,252 viral operational taxonomic units (vOTUs) (≥ 5 kb) quality and completeness (assessed with CheckV). Comp. for complete, HQ for high-quality, MQ for medium-quality, LQ for low quality, ND for not-determined. The inserted pie chart depicts an assessment of virus lifestyle (temperate, ambiguous, and NA-not assigned) of the 257,252 vOTUs (≥ 5 kb).

**Fig.S4. Pristine groundwater taxonomy assignments.**

**(a)** The percentage of virus taxonomy assignments at the class level. The complete taxonomy assignment can be seen in **Table S3**. The taxonomy assignment analysis was done using vConTACT3. **(b)** The percentage of virus taxonomy assignments at active viruses’ phylum and class level.

**Fig. S5. Active viruses and their taxonomy.**

We classify viruses as ‘active’ if at least one expressed gene per 10 kb of their genome was detected. **(a)** The heatmap depicts active viruses (transcript relative expression). Viruses are considered active if their genomes have at least one expressed gene per 10 kb. The colour intensity on the heatmap reflects the relative expression level of each virus within a specific metatranscriptome sample. Higher relative expression (in log10 scale) corresponds to more intense colours. **(b)** the proportion of active unique and shared vOTUs.

**Fig S6. Viral activity and their associated hosts.**

**(a)** The Pie charts depict the percentage of the predicted hosts of the active viruses. **(b)** The stacked-bar plots depict the viral transcripts of relative expression (%) across various categories. These categories include virus taxa lifestyle (temperate, virulent, and non-determined), host association (viruses with host links exceeding 1% abundance in Fig. 3), and those infecting DPANN, non-DPANN, CPR, and non-CPR hosts.

**Fig.S7. Groundwater vOTUs comparison across different databases**

Percentage of Hainich groundwater Protein Cluster (PCs) and virus Operation Taxonomy Units (vOTUs) that were identified in other public databases: IMG/VR v4.1^37^, Gut Virome Database (GVD, *n* = 33,242)^43^, Global Soil Virome database (GSV, *n* = 80,750)^44^, Global Oceans Viromes 2.0 database (GOV2, *n* = 195,728)^40^, Viral RefSeq v.222^45^ (*n* = 18,719), Fennoscandian Groundwater Virome (FGV, *n* = 4,051)^39^, and Rumen Virome Database (RVD, *n* = 397,180)^46^ (see materials and methods). At the protein cluster (PC), the number of the overlapping cluster and the total cluster were shown. At vOTU-level, we compared using BLASTn, and clustering at ID 95%, Cov. 50%; ID 95% and Cov. 80% (see materials and methods).

**Fig.S8. Virus comparison with IMG/VR (v.4.1) freshwater ecosystem at the vOTU-level.** The UpSet plots represent the comparison analyses done at **(a)** ≥ 95% ANI, and ≥ 80% coverage. And **(b)** ≥ 95% ANI, and ≥ 50% coverage. The intersection size represents the number of clusters.

**Fig.S9. Virus community patterns.**

**(a)** Groundwater viruses’ distribution across the well. The UpSet plot depicts the similarity and uniqueness of vOTUs across the well at ≥ 95% ANI, and ≥ 80% coverage. The colors represent well. Insert Venn diagram shows the similarity and uniqueness of vOTUs based on the oxic/anoxic wells. **(b)** vOTU relative abundance (%) is classified as a unique and shared virus community. **(c)** vOTU relative abundance (%) is classified as a core, common, and rare virus community (see materials and methods).

**Fig. S10. The ecological patterns and diversity analysis of pristine groundwater viruses.**

**(a)** Non-metric multi-dimensional scaling (NMDS). PERMANOVA was used to analyze the differences in multivariate data among viral community compositions grouped by well, filter, year, year_well, and oxic/anoxic wells. Additionally, we also used the statistical Multi-Response Permutation Procedure (MRPP) test to evaluate whether there is a significant difference between two or more predefined groups of samples based on a distance matrix. The significance delta (δ), which represents the observed within-group average distance is shown. A low δ indicates that samples within groups are more similar (less dispersed) than expected by chance. The colors represent well, and the year. And shapes represent filter size. The diversity metrics analysis, including, **(b)** Richness; **(c)** Peilous J; **(d)** Simpson; **(e)** Inverse-Simpson; and (**f**) Evenness. All pairwise comparisons shown were statistically significant (* *p*< 0.05, ** *p*< 0.01, *** *p*< 0.001, and **** *p*< 0.0001) using Kruskal-Wallis test.

**Fig. S11. The diversity analysis of pristine groundwater viruses**. **(a)** Richness; **(b)** Peilous J; **(c)**

Simpson; **(d)** Inverse-Simpson; and (**e**) Evenness for oxic and anoxic wells. **(f)** Shannons’ H; **(g)** Richness; **(h)** Peilous J; **(i)** Simpson; **(j)** Inverse-Simpson; and (**k**) Evenness based on the two years of sampling. (**l**) the Shannon’s *H* across the well on the two years of sampling. All comparisons shown were statistically significant using Wilcox test.

**Fig. S12. Microdiversity (avg. π) and GC-content (%) of across groundwater wells**. Violin plots depict

**(a)** Microdiversity (avg. π), and **(b)** GC-content (%), of unique and shared viruses (top), and core, common, and rare viruses (bottom), respectively. The Wilcoxon test was done for ‘a’, Kruskal-Wallis test for ‘b’. The statistically significant (\**p* <0.05, \*\**p* <0.01, \*\*\**p* <0.001, and **** *p*<0.0001) were shown.

**Fig.S13. Correlation between macrodiversity (Shannon’s *H*) and microdiversity (Avg. π**). Scatterplot depicts Spearman’s correlation coefficient (*R*) between macro- and microdiversity for **(a)** overall, **(b)** H14; **(c)** H32; **(d)** H41; **(e)** H43; **(f)** H51; **(g)** H52; **(h)** H53, and **(i)** based on oxic and anoxic wells. The scatterplots of **c** and **e** were analyzed with Spearman’s corr., while the rest were with Pearson’s. The number of samples per well is indicated.

**Fig.S14 Metagenomics assembled genomes (MAGs) taxonomy assignment.**

**(a)** Stacked-bar plot depicting the percentage of metagenomics assembled genomes (MAGs) recovered from Hainich groundwater. The high-(HQ) and medium-quality (MQ) MAGs. **(b)** The pie chart depicts the taxonomy assignments of MAGs, from the outer ring to the inner depicting ‘F’ family-, ‘O’ order-, ‘C’ class-, and ‘P’ phylum-level. The names of the phylum are indicated on the outer ring. The numbers in parentheses represent the count of MAGs for the respective taxa. The colors represent the phylum. **(c)** Rank abundance bar plots at phylum-level. **(d)** Phylum-level relative abundance (%) of MAGs across the groundwater (*n* = 65). The colors indicate the phylum, and only ≥1% phylum is shown. The names of the wells and size filter fractions are indicated (0.2 and 0.1 µm). **(e)** Accumulative curves of MAGs from the groundwater metagenomics (*n* = 65). The insert plot depicts the individual wells. The means are represented by colored lines, and from randomizations of samples.

**Fig. S15. The ecological patterns and diversity analysis of pristine groundwater MAGs.**

**(a)** Non-metric multi-dimensional scaling (NMDS). **(b)** Principal Coordinate Analysis (PCoA). PERMANOVA was used to analyze the differences in multivariate data among viral community compositions grouped by well, filter, year, year_well, and oxic-anoxic well. Additionally, we also used the statistical Multi-Response Permutation Procedure (MRPP) test to evaluate whether there is a significant difference between two or more predefined groups of samples based on a distance matrix. The significance delta (δ), which represents the observed within-group average distance is shown. A low δ indicates that samples within groups are more similar (less dispersed) than expected by chance. The colors represent well, and year. And shapes represent filter size. The diversity metrics analysis, including, **(c)** Shannon’s *H*; **(d)** Richness; **(e)** Peilous J; **(f)** Simpson; **(g)** Inverse-Simpson; and **(h)** Evenness across seven wells. **(i)** Shannon’s *H*; **(j)** Richness; **(k)** Peilous J; **(l)** Simpson; **(m)** Inverse-Simpson; and **(n)** Evenness across oxic and anoxic wells. (**o)** Shannon’s *H*; **(p)** Richness; **(q)** Peilous J; **(r)** Simpson; **(s)** Inverse-Simpson; and **(t)** Evenness based on the year of sampling. All pairwise comparisons shown were statistically significant (* *p*< 0.05, ** *p*< 0.01, *** *p*< 0.001, and **** *p*< 0.0001) using Kruskal-Wallis (d-h) and Wilcox test (i-t), respectively. **(u)** The relative abundance (%) of the unique and shared MAGs; **(v)** The relative abundance (%) of the core, common, and rare MAGs.

**Fig. S16. Metatranscriptome of metagenomics assembled genomes (MAGs).**

**(a)** Metatranscriptome relative expression (%) of MAGs (highest metatranscriptomic expression above 1%) across the well. (**b**) The comparison of relative expression between the oxic vs anoxic wells. Statistical analysis was done using Wilcox test. (**c**) The topmost transcribed (highest metatranscriptomic expression) genera are shown by box plots, with each point representing a single metatranscriptome (*n* = 17 metatranscriptomes). The upper and lower box edges extend from the first to third quartile, the centre line represents the median and the whiskers are 1.5× the interquartile range; points outside this range represent outliers. Only genes that are present in 10% of metatranscriptome samples (*n* = 7) were considered in the analysis. (**d**) The topmost transcribed (highest metatranscriptomic expression) genera are shown by box plots, with each point representing a single metatranscriptome (*n* = 17). The upper and lower box edges extend from the first to third quartile, the centre line represents the median and the whiskers are 1.5× the interquartile range; points outside this range represent outliers. No filtering was done for this analysis (unfiltered).

**Fig. S17. Ecological drivers’ analysis.**

Pearson’s correlation was used for analyzing environmental factors. Mantel test results were utilized to examine the relationship between environmental factors and viruses (richness, macro-, and microdiversity), as well as hosts (richness and shannon’s *H*). The figure displays only those relationships that were statistically supported.

**Fig. S18. Virus-Host links count.**

**(a)** Pie chart depicts the percentage of predicted hosts. The outer ring represents the hosts’ domains, the second ring represents the hosts’ categories (DPANN, non-DPANN, CPR, and non-CPR), and the center ring represents the hosts’ phyla. **(b)** The overall bar plots depict the count of virus-host linkages. The host links are based on the iPHop analysis (see materials and methods). Only the linkages with a confidence score ≥ 90% and those with 100 links were shown (see the rest in **Table S9**). The number of MAGs with the assigned taxonomy is shown in parentheses.

**Fig. S19. Virus-Host links distributions across the groundwater wells and virus-host correlations.**

**(a-c)** Spearman’s coefficient correlation analysis between the average relative abundance (RA), Shannon’s *H*, and Richness of viruses and hosts with links. **(d)** The Pearson’s coefficient correlation of 35 virus/host links.

**Fig. S20. The lineage-specific virus/host abundance ratio (VHRs) per well**. The lollipop plots depict the VHRs by host lineage (log_10_). The red line indicates the 1/1 VHR. The host rank abundance is depicted as a grey bar plot.

**Fig. S21. Virus-host links count.**

**(a)** Categorized as DPANN, non-DPANN, CPR, and non-CPR. Only the linkages with a confidence score ≥ 90% and those with 100 links were shown (see the rest in **Table S8**). The numbers in parentheses ‘()’ represent the number of MAGs. While (REF) means link to genomes in iPHop references. The colors represent taxonomy assignment and taxa-category. **(b-c)** Phylum-level relative abundance (%) for each sample (*n* = 65) for the predicted host (top) and viruses (bottom). Categorized as DPANN, non-DPANN, CPR, and non-CPR. **(d)** The bar plot depicts the number of virus links to CPR classes and **(e)** to DPANN.

**Fig. S22.** Co-occurrence network to assess the potential association of CPR and DPANN with other bacterial and archaeal MAGs for 2022 sampling year. The circles depict the potential bacteria hosts of CPR and DPANN. The red diamonds represent CPR, and the dark-grey triangle is for DPANN. The zoomed-in clusters highlight some associations of CPR and DPANN MAGs with potential host MAGs, e.g. Chloroflexota, and Proteobacteria. Virus links in the inserted figure were identified through *in silico* host prediction analysis. Pie charts illustrate the percentage of MAG (*n* = 1,275) with CPR/DPANN associations based on the co-occurrence network analysis.

**Fig. S23. Prophage identified in MAGs.**

**(a)** The workflow to identify prophages from 1,275 MAGs. **(b)** The percentage of MAGs with and without identified prophages. **(c)** The barplot depicts the number of prophage (599 vOTUs) identified across the MAGs. **(d)** The quality assessment of prophage (599 vOTUs) genome quality via CheckV. **(e)** Comparison between vOTUs identified in metgenomics data (metaG) and those from MAGs. The lustering analysis was done using a threshold of 95% identity and 80% coverage. About 94.5% vOTU prophages overlapped with those found in metagenome data. **(f)** Phylum-rank taxonomy of the MAGs with the overlapping vOTU prophage.

**Fig. S24. Putative auxiliary metabolic genes (AMGs) targeted some hosts’ metabolisms.**

**(a)** Some examples of viruses-carrying AMGs. The color legend indicating the AMGs, virus genes, bacterial genes, and hypothetical genes is shown. **(b)** sulfur metabolism, **(c)** nitrogen metabolism, **(d)** methane and related metabolisms, and **(e)** selected carbon metabolism. The putative AMGs are indicated in italics and KO IDs in parentheses. The AMGs and the metabolic process that viruses target are colored in red.

**Fig. S25 Example of the expression of AMGs across the groundwater.**

The heatmap depicts normalised transcript coverages (relative expression) for genes affiliated with each pathway.

**Fig. S26. Identified putative AMG in CPR and DPANN.**

**(a)** The pie charts illustrate the AMGs identified in CPR. **(b)** The pie charts illustrate the AMGs identified in DPANN. The colors indicate the AMG categories and further descriptions.

## Supplementary tables

**Supplementary Table 1**. Sampling and metagenomics data overview

**Supplementary Table 2**. Virus overview: quality assessment, lifestyle

**Supplementary Table 3**. Virus taxonomy assessment based on vConTACT3

**Supplementary Table 4**. Normalized relative expression of viruses, scaling factor for metatranscriptome normalization, and statistical comparison

**Supplementary Table 5**. Normalized relative abundance table for vOTUs and MAGs

**Supplementary Table 6**. Virus diversity assessment

**Supplementary Table 7**. Groundwater environmental data

**Supplementary Table 8**. Virus-host link, MAGs used in the analysis, MAGs-virus link, and MAGs metabolic capabilities

**Supplementary Table 9**. Virus-host ratio (VHR) and comparison analysis across the well

**Supplementary Table 10**. Identified CRISPR-Cas spacers in the groundwater MAGs, Spacers’ BLATST analysis (at different mismatch cutoffs) and IMG/VR blast

**Supplementary Table 11**. CPR co-occurrence analysis

**Supplementary Table 12**. Auxiliary metabolic genes (AMGs) list

**Supplementary Table 13**. Metabolic reconstruction for host pathways and AMG

